# Improving COVID-19 Testing Efficiency using Guided Agglomerative Sampling

**DOI:** 10.1101/2020.04.13.039792

**Authors:** Fayyaz Minhas, Dimitris Grammatopoulos, Lawrence Young, Imran Amin, David Snead, Neil Anderson, Asa Ben-Hur, Nasir Rajpoot

## Abstract

One of the challenges in the current COVID-19 crisis is the time and cost of performing tests especially for large-scale population surveillance. Since, the probability of testing positive in large population studies is expected to be small (<15%), therefore, most of the test outcomes will be negative. Here, we propose the use of agglomerative sampling which can prune out multiple negative cases in a single test by intelligently combining samples from different individuals. The proposed scheme builds on the assumption that samples from the population may not be independent of each other. Our simulation results show that the proposed sampling strategy can significantly increase testing capacity under resource constraints: on average, a saving of ~40% tests can be expected assuming a positive test probability of 10% across the given samples. The proposed scheme can also be used in conjunction with heuristic or Machine Learning guided clustering for improving the efficiency of large-scale testing further. The code for generating the simulation results for this work is available here: https://github.com/foxtrotmike/AS.

## 1. Introduction

Effective large scale testing and contact tracing have been successfully used in a number of countries for controlling the spread of the SARS-CoV-2 virus (CoV-2) which causes COVID-19 disease [1]. However, in resource-limited settings, it may not be feasible to do large scale testing unless the efficiency of existing tests is improved in terms of number of tests required for a given number of samples. In this short paper, we discuss a computer-science inspired divide and conquer strategy based on pooling samples from multiple individuals that can improve test efficiency by a significant amount under a minimalistic set of assumptions.

## 2. Methods

### 2.1 Assumptions

Given a set of *N* individuals to be tested for CoV-2, the number of tests *T* required for identifying positive individuals can be reduced from *N* by considering the fact that the probability of testing positive *p* is small (say, *p* = 0.1) and individual test results are typically not independent of each other (e.g., members in the same household or people in contact with each other or other CoV-2 infected individuals can have dependencies in their test results). In this work, we propose a divide and conquer agglomerative sampling strategy that is built on these ideas and can be used to reduce the number of tests. Before delving into the details of the method, we present a list of assumptions underlying the proposed method:

1. Pooling: Samples of multiple individuals can be combined or mixed into a single “bag” which can be tested by a single test such that:
  a. The test produces a positive outcome if any of the samples in the bag is positive
  b. The test produces negative outcome if none of the samples in the bag is positive
  c. Testing in bags does not change the error rate of the test being used
2. Multiplicity: Multiple samples can be taken from a single individual or a single sample from an individual can be divided further
3. Rarity: The probability of testing positive is small (*p* < 0.2)

It can be expected that these assumptions are satisfied by a number of current tests for CoV-2 infection such as the quantitative RT-PCR and serological (antibody) testing [2].

### 2.2 Algorithm Description

Consider a set of *N* individuals *S* = {1,2, …, *N*} to be tested for CoV-2 infection. Assume that the (originally unknown) test result of each of the individuals is given by *y*_*i*_ ⊂ {0,1}, *i* = 1 … *N*. Without loss of generality or introducing any limitations in the model, assume that for each individual, we are also given a set of **d**-features *x*_*i*_ ∈ *R*^*d*^ (such as frailty, age, gender, contact with known or suspected CoV-2 infected patients, geographical location, symptoms, family/household dependencies, etc.,) that can be used to generate a degree of belief of that individual to test positive. We denoted this degree of belief by *b*_*i*_, *i* = 1 … *N*. In case, it is not possible to assign a belief to each individual, *b*_*i*_ can be considered to be uniformly random, i.e., *b*_*i*_~*U*(0,1). Alternatively, belief can be assigned by a human oracle in a subjective manner or can be obtained through machine learning or probabilistic modelling based on the given set of features. If we cluster or mix individual samples into bags and proceed with testing these bags in a hierarchical manner, the number of required tests can be reduced by essentially pruning out multiple negative samples in a single test. For this purpose, consider a tree structure organization of the given set of individuals based on the degree of belief *b*_*i*_, *i* = 1 …*N* (or using the given set of features directly) as shown in the example figure below. The basic idea of agglomerative testing is that we test a bag of samples and if the bag level result comes out negative, then there is no need to test each of the samples individually. However, in case, the test comes out positive, we subdivide the samples into further clusters and test each of these bags next. This is continued until we get a test score of each individual. Furthermore, if a test for a bag comes out positive but the next sub-bag tests negative, then we know that the positive result is a consequence of a positive individual in the other bag which can be split further directly without additional testing. This guide algorithm based on even binary split is summarized in Algorithm-1. The figure below shows that if we obtain a mixed bag of all individual samples 1-8 and do a single test, the outcome will be negative and there is no need to do individual testing. For a bag comprising of cases 9-16, the result of the test will be positive because there is at least one positive individual in the bag. Doing this in a recursive manner can lead to reducing the number of tests required from 16 to 11 or 14 depending upon how the terminal nodes are tested.

If we have access to informed belief values, then the given samples can be sorted with respect to their belief values prior to tree construction. Tree construction can also be done in an unsupervised manner based on existing individual features coupled with hierarchical or agglomerative clustering. [3].

### 2.3 Simulation Setup

In order to evaluate the efficacy of this approach, we constructed a simple simulation in which *N* individuals are assigned random test labels (*y*_*i*_ = 1 with probability *p* and *y*_*i*_ = 0 with probability 1 − *p*). Each individual is then assigned a degree of belief *b*_*i*_. We tested with both a random degree of belief (no belief information) and varying degrees of belief as measured by the concordance between *b*_*i*_ and *y*_*i*_ by using an additive normal distribution noise prior *b*_*i*_ = *y*_*i*_ + *n*_*i*_ (with *n*_*i*_~ℵ(0,*σ*)) with the degree of noise controlled by the standard deviation parameter *σ*. For a given simulation setting (number of individuals, prior probability and belief control factor *σ*), we calculate the number of required tests *T* by the proposed sampling method. In order to get reliable statistical estimates of the distribution of the number of required tests for a given simulation setting, we repeated the simulation multiple times with the same setting and plotted the distribution of the number of required tests using a box plot.

#### Algorithm 1 The proposed method for Agglomerative Sampling

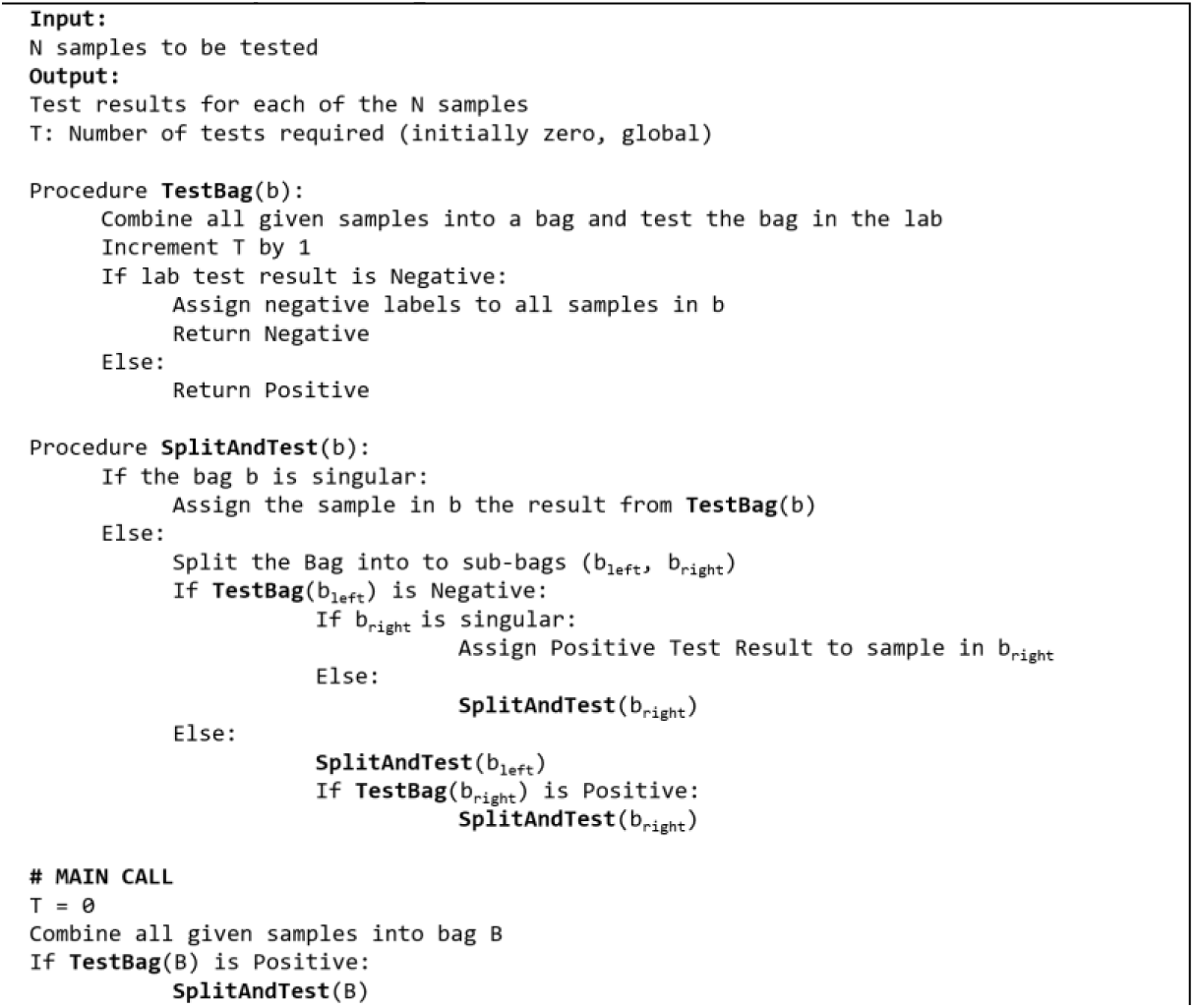

**Figure 1.**
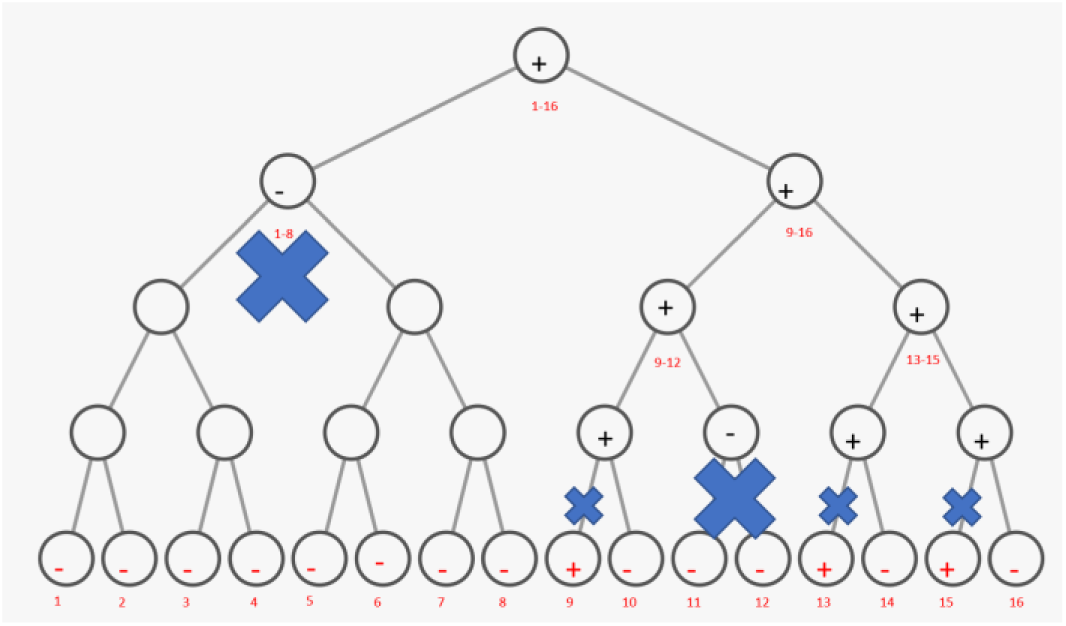
Concept diagram for the proposed agglomerative sampling scheme for a given set of 16 individuals with 3 positives (p=0.19) indicated by the plus (+) or (-) sign in the leaf nodes of the tree. Each circle represents a possible test of a bag of samples. Each x indicates pruned nodes. Note that the number of tests required is 11 instead of 16.

### 2.4 Mathematical Analysis

Based on our computational analysis, the expected number of tests required for a given positive probability and input samples (under no belief assumptions) can be calculated as: *t*(*p*,*N*) = 2(*N* − 1)(1 − 2^−4.5*p*^) + 1. This formulation captures the typical average case number of required tests using the proposed strategy. It can be seen that this formulation is heavily dependent on the value of the positive probability. However, it can significantly reduce the number of required tests when the probability is small, e.g., for community level testing. The probability value up to which the proposed strategy can remain effective, i.e., up till T<N, is called the utility breakdown probability *p*_*d*_ = *p*|_*t*(*p*,*N*)<*N*_ and it is independent of the value of *N* (for large N) can be calculated as: *p*_*d*_ = 0.22. (proof omitted for brevity).

### 2.5 Lab Testing

Lab testing of the proposed method is currently underway. However, we are sharing the basic idea of the proposed method together with the simulation results in order to support the ongoing COVID-19 efforts across the globe. Specifically, our planned wet lab experiments will be aimed at studying the impact of this approach on the sensitivity/sepecificity of tests and understanding practical limitations for use with PCR or immunoassay-based testing as well as serology.

## 3. Results

### 3.1 Under no informed heuristic or belief

All simulation results guarantee that the output of the tests remains unchanged from individual testing, i.e., if the original test identifies a given sample as positive (negative) then using the proposed scheme will identify that given sample as positive (negative) but with fewer number of tests required in overall. Figure 2 shows the number tests required under the proposed scheme for different positive probability values (*p*) and different values of (*N*) under no a prior belief (uniformly random *b*_*i*_). It can be clearly seen that the mathematical formula for the number of expected tests is in excellent concordance with the simulation results. Figure 2(a) shows that the number of required tests for *N* = 16 with the proposed method remains below *N* up to a probability of *p*_*d*_ = 0.22 as expected. Figure 2(b) shows the same analysis for *N* = 256. Figure 2(c) shows that the expected number of tests that can be saved is above 40% for all values of *N* at a positive probability of *p* = 0.1. This clearly shows that the proposed scheme can be very beneficial in practical settings. The number of required tests can be reduced further by incorporating a belief parameter or performing unsupervised agglomeration based on individual features as discussed below.

**Figure 2.**
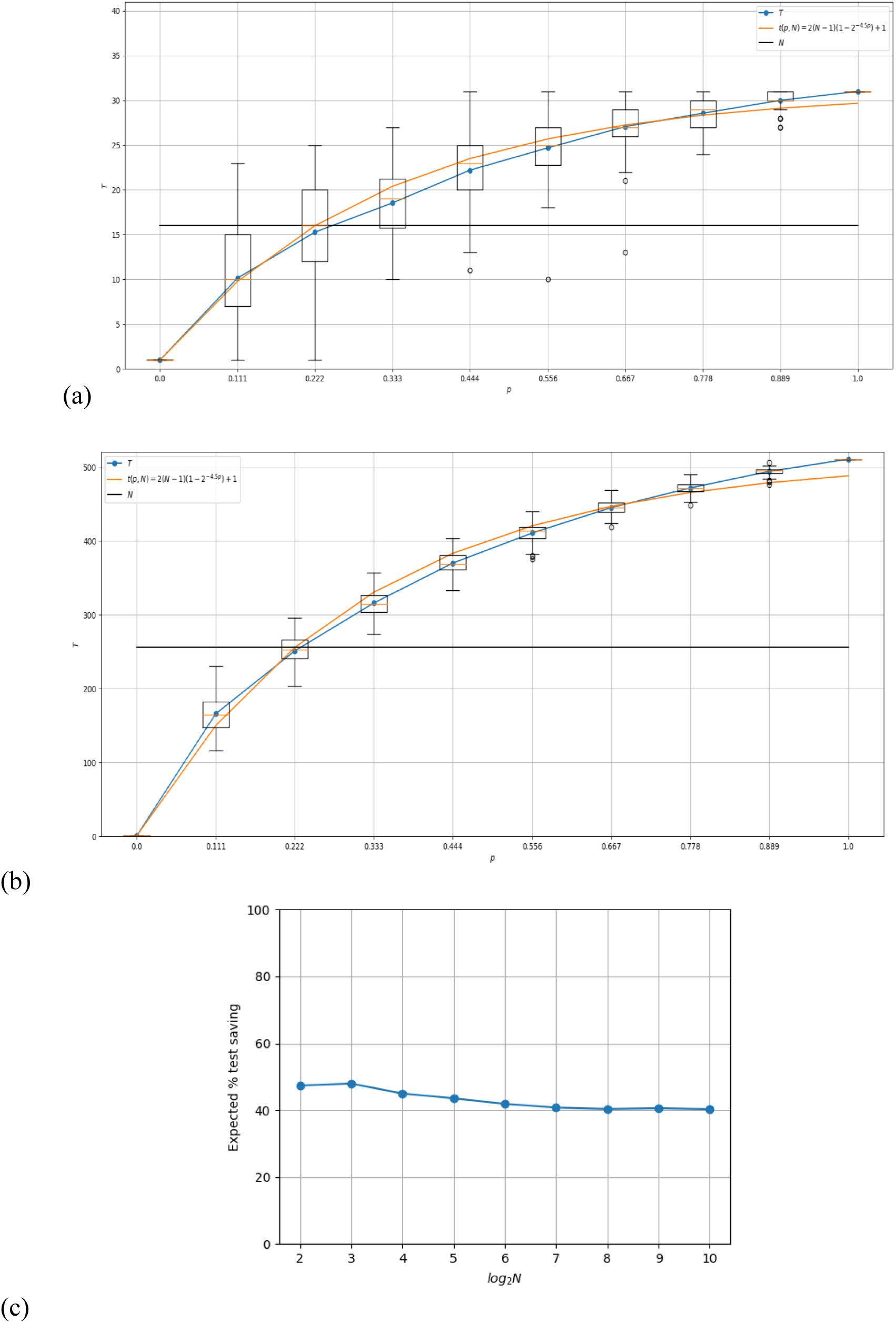
Simulation results of the proposed sampling scheme. In each figure, a plot of the average number of tests required in multiple trials for a given positive probability value are shown as a box plot together with the theoretical estimate: t(p,N) = 2(N − 1)(1 − 2^−4.5p^) + 1.(a) for N=16 (b) for N = 256 With σ = 1.0 and (c) Plot of the number of tests saved as a function of the N for p=0.1.

**Figure 3.**
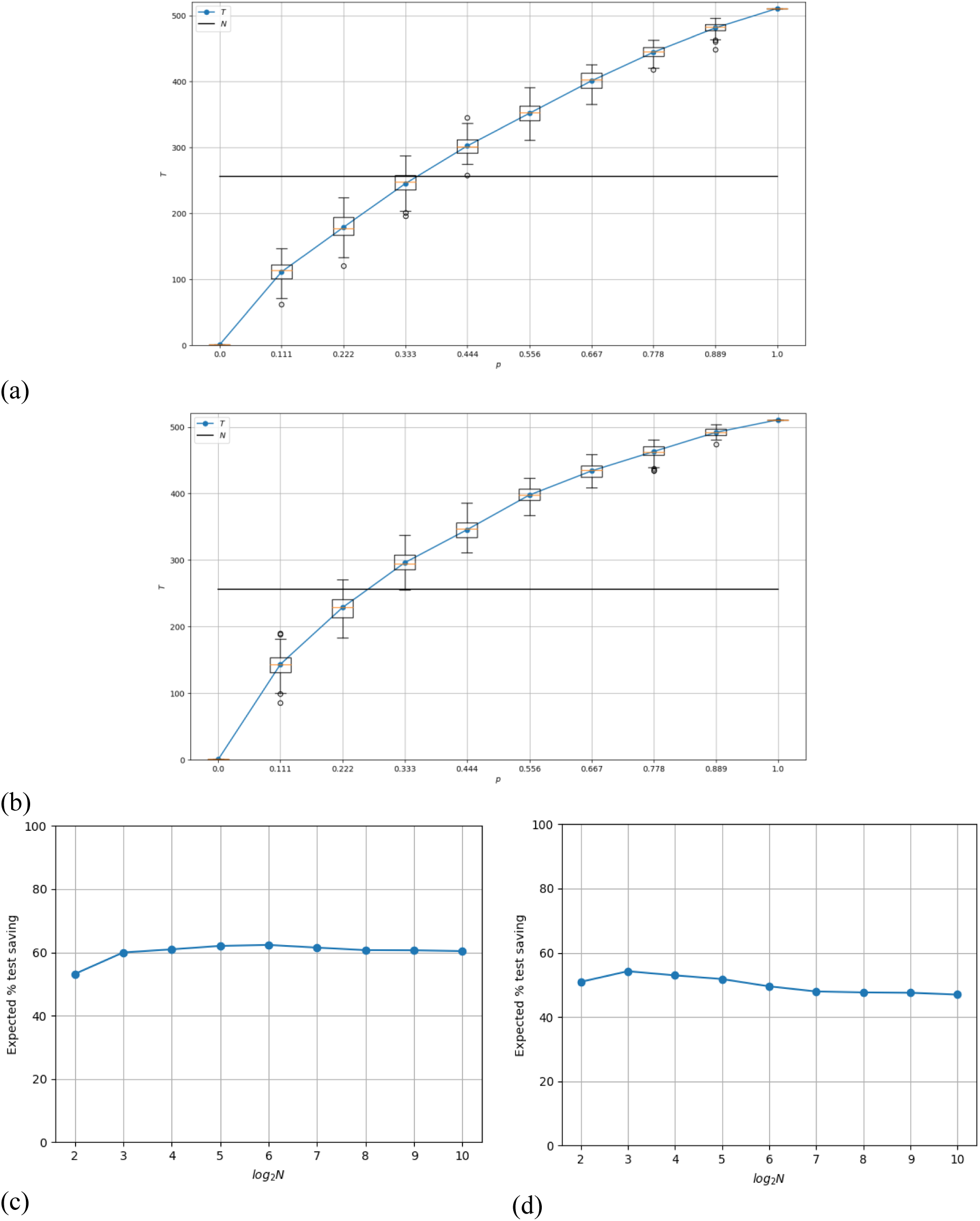
Results of the simulation in terms of number of tests required for different positive probability values for N=256 with (a) σ = 0.5 and (b) σ = 1.0. (c) and (d) show the expected percentage tests saved for different values of N for p=0.1 with σ = 0.5 and = 1.0, respectively.

### 3.2 Under an informed heuristic or belief

As discussed in the methods section, if there is a way of predicting the likelihood of someone testing positive for CoV-2 (e.g., by using a machine learning method) or assigning such belief based on expert opinion, then the efficiency of the proposed scheme can be further improved by first ranking (sorting) the given samples with respect to their belief values. The concordance of the belief value *b*_*i*_ and the true status *y*_*i*_ can be measured by using the area under the receiver operating characteristic curve (AUC) between these values [4]: *AUC* = 0.5 implies poor concordance between belief and the actual test status whereas *AUC* = 1.0 implies perfect concordance. Please note that this AUC score is not between the test outcomes and the actual status but is used as a means of measuring the impact of the additive noise on the belief values for each individual. The degree of concordance is dependent upon the value of the noise factor: *σ*: *σ* = 0 will result in perfect concordance (*AUC* = 1) in which case, no testing is needed as the belief is perfect whereas for large values of *σ*, the AUC value will be 0.5. Below we show the results of the proposed scheme for various values of *N*,*p* and *σ*. For *σ* = 1.0, we get an average AUC score of 0.75 and this leads to a moderate increase in the number of tests that can be saved in comparison to the no-belief simulation. This shows that even a weak belief assignment model coupled with the proposed scheme can significantly reduce the number of required tests. For *σ* = 0.5 (with an AUC score of 0.9), the saving is even more substantial (up to 60%). This clearly shows that the proposed testing scheme can lead to further improvements by incorporating belief through machine learning models or expert assignment.

## 4. Conclusions and Future Work

In this work, we have developed a community laboratory testing strategy for CoV-2 based on a divide and conquer approach [5] that can reduce the number of tests required for testing a given number of samples. It can optionally be used in conjunction with a belief assignment method such as a machine learning prediction model or with guidance from a human expert to improve testing efficiency even further. In terms of machine learning, the proposed scheme can be adapted for use to work together with a machine learning model which generates a ranked list of likely positive samples which can be tested individually followed by agglomerative testing of the remaining samples. Additionally, in the absence of a predictive model or another means of belief assignment, the proposed scheme can use feature-based unsupervised clustering to reduce the number of required tests building on the assumption that the test results of individuals are not independent of each other.

We have opted to share the proposed method in the hope that it can be beneficial to large-scale CoV-2 testing and the management of patients with COVID-19. Laboratory trials with the proposed sampling technique are being considered at the University of Warwick to study the impact of the proposed strategy on accuracy of existing testing methods and understand practical limitations.

## References

[1] Hadaya, Joseph, Max Schumm, and Edward H. Livingston. “Testing Individuals for Coronavirus Disease 2019 (COVID-19).” JAMA, April 1, 2020. https://doi.org/10.1001/jama.2020.5388.

[2] Hogan, Catherine A., Malaya K. Sahoo, and Benjamin A. Pinsky. “Sample Pooling as a Strategy to Detect Community Transmission of SARS-CoV-2.” JAMA, April 6, 2020. https://doi.org/10.1001/jama.2020.5445.

[3] Aggarwal, Charu C., and Chandan K. Reddy. Data Clustering: Algorithms and Applications. CRC Press, 2013.

[4] Alpaydin, Ethem. Introduction to Machine Learning. Cambridge, Mass.: MIT Press, 2010.

[5] Cormen, Thomas H., Charles E. Leiserson, Ronald L. Rivest, and Clifford Stein. Introduction to Algorithms. 3rd edition. Cambridge, Mass: MIT Press, 2009.

